# Unusual Features of the Membrane Kinome of *Trypanosoma brucei*

**DOI:** 10.1101/2020.12.08.416529

**Authors:** Bryan C. Jensen, Pashmi Vaney, John Flaspohler, Isabelle Coppens, Marilyn Parsons

## Abstract

In many eukaryotes, multiple protein kinases are situated in the plasma membrane where they respond to extracellular ligands. Ligand binding elicits a signal that is transmitted across the membrane, leading to activation of the cytosolic kinase domain. Humans have over 100 receptor protein kinases. In contrast, our search of the *Trypanosoma brucei* kinome showed that there were only ten protein kinases with predicted transmembrane domains, and unlike other eukaryotic transmembrane kinases, seven are predicted to bear multiple transmembrane domains. Most of the ten kinases, including their transmembrane domains, are conserved in both *Trypanosoma cruzi* and *Leishmania* species. Several possess accessory domains, such as Kelch, nucleotide cyclase, and forkhead-associated domains. Surprisingly, two contain multiple regions with predicted structural similarity to domains in bacterial signaling proteins. A few of the protein kinases have previously been localized to subcellular structures such as endosomes or lipid bodies. We examine here the localization of epitope-tagged versions of seven of the predicted transmembrane kinases in *T. brucei* bloodstream forms and show that five localized to the endoplasmic reticulum. The last two kinases are integral membrane proteins associated with the flagellum, flagellar pocket, or adjacent structures, as shown by both fluorescence and immunoelectron microscopy. Thus, these kinases are positioned in structures suggesting participation in signal transduction from the external environment.

## Introduction

With its complex life cycle spanning different hosts and tissues, the response of *Trypanosoma brucei* to its environment is a subject of considerable interest. In other eukaryotes, signaling by extracellular molecules is initiated by ligand binding, typically to proteins situated at the plasma membrane. The most prominent categories of such proteins are G-protein coupled receptors (absent in trypanosomatids) and protein kinases (PKs) [1]. *T. brucei* also possesses a large family of surface proteins that have external ligand-binding domains and internal adenylate cyclase domains [2], although the ligands and signaling pathways involved remain to be elucidated. Transporters can also function in signal transduction by internalizing regulatory molecules such as Pi or cGMP [3-5]. In *T. brucei*, a plasma membrane carboxylate transporter is important in regulating the transition of mammalian bloodstream forms (BF) to the insect procyclic form (PF) stage [6].

Eukaryotes tend to have hundreds of PKs that bear the signature eukaryotic PK (ePK) motifs in the catalytic domain, *T. brucei* has ∼180 protein-coding regions that bear a canonical ePK domain [7]. Almost all PK orthologues are shared among *T. brucei, Trypanosoma cruzi* and *Leishmania major*. Most eukaryotes have tens to hundreds of PKs that bear transmembrane domains (TMDs), and in metazoans almost all are tyrosine kinases that reside in the plasma membrane. Ligand binding to the extracellular domain generates a signal that is transduced to the cytosolic catalytic domain resulting in kinase activation. In a previous analysis of the genome, we identified ten PKs predicted to bear TMDs as predicted by TMHMM [7], The redrawing of CDS boundaries based on 5’ transcript mapping eliminated the putative TMD of one (Tb927.10.16160) and annotation of another PK was updated to include a TMD (Tb927.10.1910). The catalytic domains of these PKs cluster with serine/threonine kinases [7].

About half of the *T. brucei* TMD PKs have been characterized functionally. Tb927.2.2720, dubbed *MEKK1* (so-named for the similarity of its catalytic domain to MAP kinase kinase kinases), is required for the quorum sensing pathway that triggers proliferating slender BF to enter stationary phase as a prelude to transmission [8]. Tb927.11.14070, known as *RDK1*, encodes a repressor of BF to PF differentiation [9]. Tb927.11.8940 encodes LDK (lipid droplet kinase), which is present on lipid droplets and essential for their biogenesis [10]. Tb927.4.2500, which encodes an eIF2 kinase-related protein, EIF2K2K, is localized to the flagellar pocket [11]. While the function of this kinase in *T. brucei* is not known, its orthologue in *Leishmania* appears to regulate promastigote to amastigote development [12] and the *T. cruzi* orthologue similarly regulates the development of infective epimastigote forms [13]. Tb927.10.1910 is suggested to play a role in melarsoprol sensitivity on the basis of high-throughput RNAi assays [14]. Cell death was not observed upon depletion of the TM kinases in BF in a kinome-wide stem-loop RNAi study, although the degree of knockdown was not quantitated [9].

Here we report studies examining the TMD PKs. Many of them have unusual domain structures, such as the presence of multiple TMDs, nucleotide cyclase domains, or regions of structural similarity to bacterial signaling modules. We examined the subcellular localization of seven of these PKs in monomorphic BF, which are unable to differentiate to PF. Five of the tagged PKs localized predominantly to the endoplasmic reticulum (ER), indicating they entered the secretory system. Two were found to have particularly distinctive distributions: FHK localized to a region at or near the flagellar pocket and MEKK1 localized similarly, as well as the flagellum. Immunoelectron microscopy of cultured PF confirmed the unique positioning of these two kinases.

## Results

### Domain structure

Figure 1A depicts schematics of the ten *T. brucei* PKs that have TMDs, showing the location of the TMDs. Also shown are the PK catalytic domains and additional domains listed on TriTrypDB (based primarily on sequence similarities), plus regions HHpred identified as having predicted structural similarities to other protein modules. These PKs are large, with the smallest (the lipid droplet kinase LDK) being 553 aa, about twice the size of the PK domain, and the rest ranging from 929 to 1678 aa. The PK domain of all of these kinases, with the exception of LDK, resides at the C-terminus.

**Figure 1.**
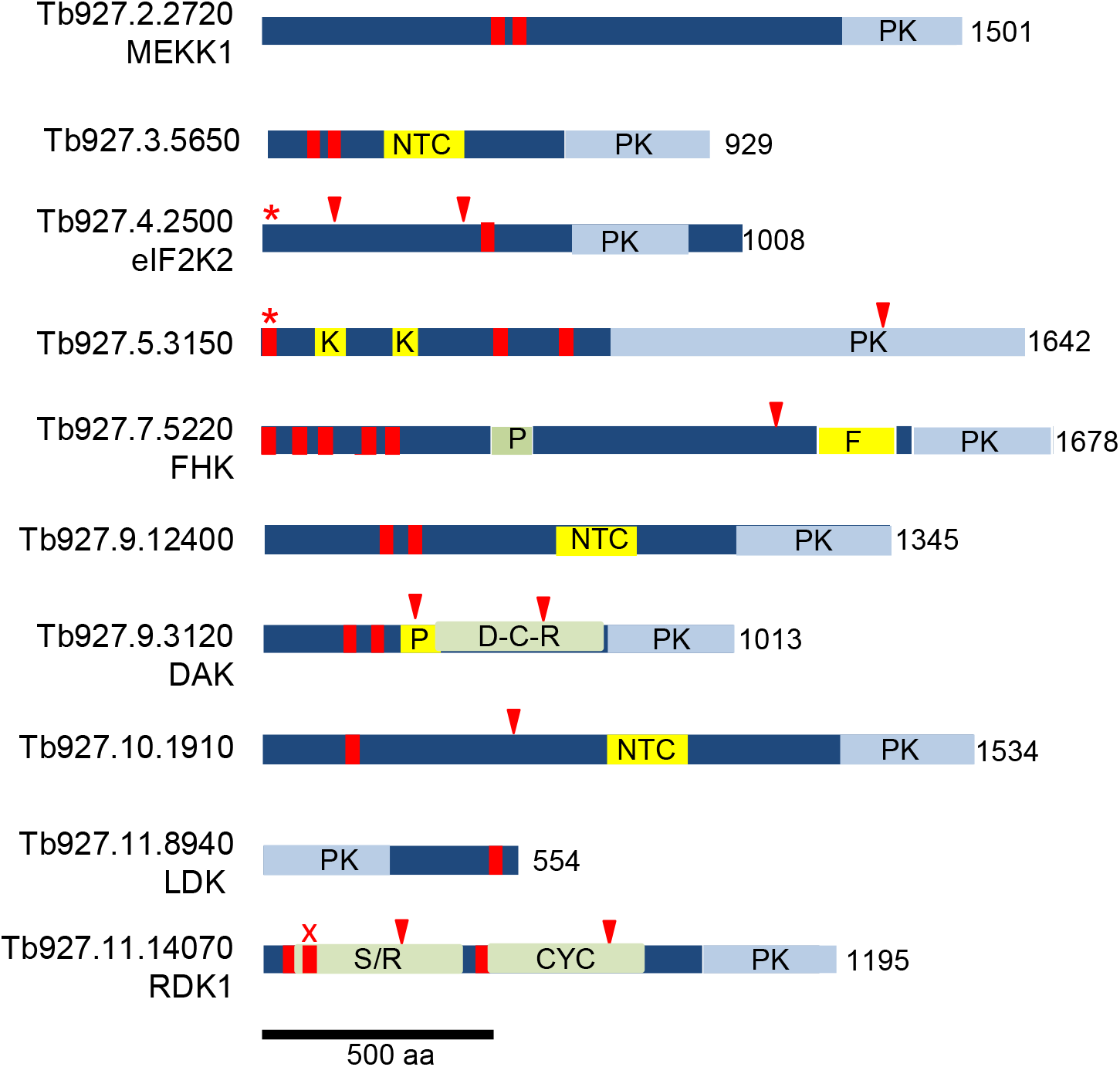
Schematics of *T. brucei* protein kinases predicted to bear transmembrane domains. TMDs predicted by both TMHMM and CCTOP are marked with red bars (one predicted by TMHMM only is marked with a red X); additional TMDs predicted by CCTOP are marked by red arrowheads. Predicted signal sequences are indicated by a red asterisk. Light blue regions are ePK Pfam domains. Yellow bars depict other domains: PAS motif (P), forkhead associated domain (F), nucleotide cyclase-like domain (NTC), Kelch propeller blade (K). Light green bars demarcate regions with predicted structural similarity to known structures revealed by HHpred. Bacterial DHp, catalytic, and receiver domains characteristic of histidine kinases (D-C-R) in Tb927.9.3120, as well as bacterial sensor/receptor domains (S/R) and adenylyl/guanylyl cyclases (AGC) in RDK1, exhibited probabilities >95% (see Figs. 2 and S2 for details). The PAS-like domain in FHK was detected with >90% probability. Length of the predicted protein in amino acids is indicated.

#### Transmembrane domains and signal peptides

Surprisingly, seven of the ten proteins are predicted by TMHMM to be multi-pass proteins, having two or more TMDs, a topology that is extremely uncommon for eukaryotic PKs. We therefore re-examined the TMD (predicted by TMHMM and shown TriTrypDB, red bars in Fig. 1) using CCTOP (Constrained Consensus TOPology Prediction, http://cctop.enzim.ttk.mta.hu/, which combines ten algorithms with structural data to yield a model. The CCTOP models concurred with TMHMM for four PKs and proposed additional TMDs for the rest of the PKs (Fig. 1, red arrowheads), and rejected one of the TMDs on RDK1 (red x). Some of the additional TMDs predicted with CCTOP are unlikely to be authentic since they occur in the middle of domains, for example, one that punctuates the ePK catalytic domain in Tb927.5.3150, and those within structural domains of RDK1 and Tb927.9.3120. When we examined the orthologues of these *T. brucei* PKs in *L. major* and *T. cruzi*, the number and position of the TMHMM-predicted TMDs generally concurred (Table S1). The exceptions included the *L. major* orthologue of eIF2K2 (LmjF.34.2150), the *T. cruzi* CL-Brener orthologues of Tb927.5.3150, and the *T. cruzi* orthologue of LDK (TcCLB.511801.14), all of which lack a TMHMM-predicted TMD. However, CCTOP did predict TMDs in these proteins. Furthermore, since LDK resides in a monolayer membrane surrounding lipid droplets rather than a bilayer, algorithms may be less proficient at identifying relevant membrane insertion/association sequences. While it is not possible to be certain of the number of TMDs from bioinformatic predictions, an even number of TMDs would place the N-terminal region and C-terminal catalytic domain in the same compartment. For example, MEKK1 is predicted to have a hairpin of two TMDs, separated by only four aa. The extended N- and C-regions would either both be cytosolic, or both be extracellular. We are unaware of any transmembrane kinases that display catalytic domains on the external face of the plasma membrane, so propose that the former possibility is most likely.

Most membrane proteins destined for the secretory system or plasma membrane enter the ER membrane co-translationally, courtesy of a signal peptide or a transmembrane domain [15]. Interestingly, it has been proposed that two TMDs separated by loop, such as seen in MEKK1 and several other TMD kinases (Fig. 1), may be a topogenic signal for insertion into the ER membrane [16]. Two of the PKs bearing TMDs are also predicted to bear signal peptides: eIF2K2 and Tb927.5.3150. Three additional *T. brucei* PKs are annotated as bearing signal peptides, but they lack predicted TMDs.

#### Domains identified by sequence similarity

Searches based on sequence similarity showed that Tb927.5.3150 bears two Kelch/Kelch-like domains, which often participate in protein-protein interactions. Additional domains related to signaling functions were detected on several PKs. Three bear nucleotide cyclase-like domains (Tb927.3.5650, Tb927.9.12400, and Tb927.10.1910). Interestingly these are the only *T. brucei* PKs identified as having such domains, although the parasite genome encodes multiple adenylyl cyclases with cytoplasmic catalytic domains and presumed ligand-binding extracellular domains [17]. One PK (Tb927.9.3120) contains a region related to PAS (Per-Arnt-Sim) domains, which usually function to bind small molecules or proteins intracellularly. Forkhead kinase, FHK, Tb927.7.5220) is so-named for the presence of a forkhead-associated domain; these domains typically interact with phosphopeptides (usually phosphothreonine in specific contexts) [18].

#### Conserved folds identified by structural predictions

The PKs were also subjected to HHpred analysis to identify regions with predicted structural similarity to protein structures available in the protein database PDB [19] (Fig. 1). The domains mentioned above all showed numerous matches with the related domains on other proteins, with probability scores of over 99%. A PAS-like region on FHK was additionally identified by HHpred, with multiple hits having a probability above 90%.

Surprisingly, two PKs have large regions of predicted structural similarity to molecular signaling modules present in bacteria. RDK1 includes two regions with predicted structural similarity to other signaling molecules (Fig. S1). The first is similar to bacterial sensor/receptor domains, such the ligand binding domain of *Pseudomonas aeruginosa* histamine receptor TlpQ [20] and the extracellular domains of a set of bacterial histidine kinases for which the ligands are unknown [21]. This region is followed by a predicted TMD and then a segment predicted to fold similarly to full-length catalytic domains of several bacterial and eukaryotic adenylyl/guanylyl cyclases. The cyclase region is followed by the eukaryotic PK catalytic domain. This organization suggests that the sensor-like domain lies on the opposite side of the membrane to the cyclase and PK domains. The modular organization of RDK1 is conserved in other trypanosomatids, as well as the more distantly related free-living bodonid *Bodo saltans* [22].

Tb927.9.3120 has co-linear regions predicted to fold similarly to molecules involved in bacterial signaling: histidine kinases and response regulators. The histidine kinases typically contain sensing modules, followed by DHp dimerization regions, then catalytic/ATP binding domains (which phosphorylate a histidine on DHp). Response regulators include receiver domains (which receive the phosphate from DHp) and effector domains. In some cases, the kinase and receiver reside on the same molecule (hybrid histidine kinases), although two component systems are more common. Importantly, in Tb927.9.3120 the regions corresponding to each domain are essentially complete and ordered as in hybrid bacterial histidine kinases [23, 24] (see Fig. 2). This organization of a hybrid histidine kinase cassette followed by a eukaryotic protein kinase domain is conserved in the Tb927.9.3120 orthologues in other trypanosomatids, such as *T. cruzi* and *L. major*, and the bodonid *Bodo saltans* (data not shown). Together, this provides intriguing evidence of a bacterial genetic incursion of a signaling cassette into a common ancestor of bodonids and trypanosomatids. Accordingly, we propose to name this gene dual ancestor kinase (*DAK*).

**Fig. 2.**
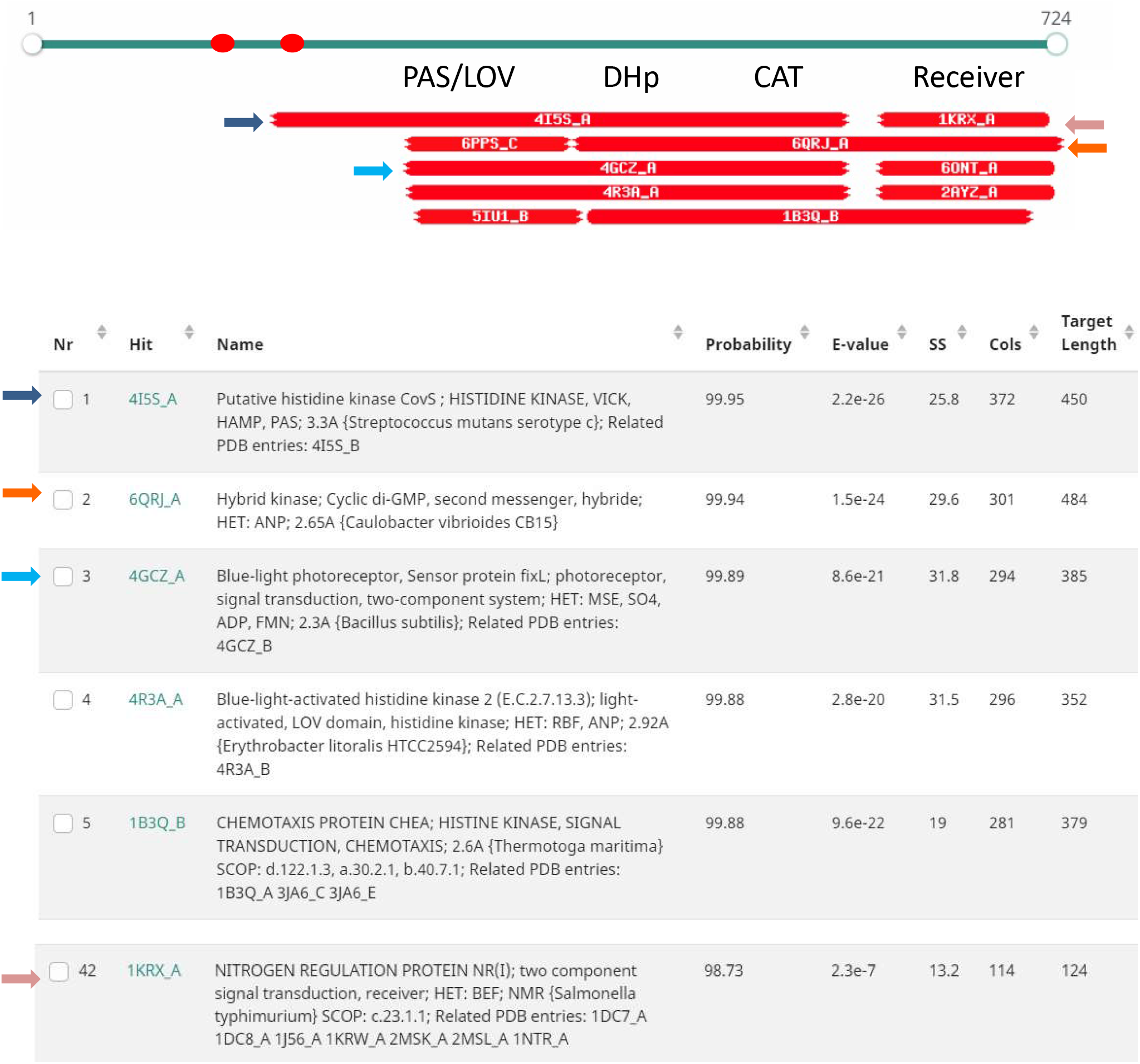
HHpred analysis of Tb927.9.3120 shows evidence of ancient domains. The PK domain (aa 737-980) is not shown. Red ovals – transmembrane domains predicted by TMHMM and CCTOP. Each bar below the gene model represents the top hits by HHpred. The color of the arrows mark descriptions matching the hits. The description includes the PDB number and chain designation (Hit), a description of the PDB entry including related PDB entries (Name), the probability of the hit based on the Hidden Markov Model (Probability), the probability of the match in an unrelated database (E-value), score for the secondary structure prediction (SS), number of amino acids aligned (Col), and the total length of the target in PDB (Target Length). The documentation for HHpred considers “Probability” the most important criterion with our hits meeting at least one of the twomost stringent criteria: having a score >95% or having a score >50% and making reasonable biological sense [19].

### Expression

We examined the expression of these PKs using our previous genome-wide ribosome profiling data from pleiomorphic slender form BF (isolated from mice) and cultured PF [25]. Ribosome profiling provides a direct measure of protein production. In Fig. 3, the dark blue and red bars show the abundance of ribosome-associated mRNA footprints of BF and PF respectively. For comparison, the mRNA abundances are also shown (light blue and pink bars). Several of the PKs show higher protein production in slender BF than in PF, with MEKK1 showing 3-fold more expression in slender BF and eIF2Ka having 2-fold more. However, the most dramatic difference was for RDK1 with 7-fold more mRNA and 40-fold more protein production in slender BF than in PF. Conversely, Tb927.9.3120 had about twice as much protein production in PF than slender BF, although the mRNA levels are very similar between stages. Most of the PKs were close to the median level of protein production for all non-pseudogenes (224 reads per kb in BF and 202 in PF). However, three PKs showed lower protein production, being in the lowest quartile of all PKs. LDK protein production was low in both stages; FHK protein production was low in BF and very low in PF; and Tb927.9.12400 protein production was almost absent in BF and very low in PF.

**Fig. 3.**
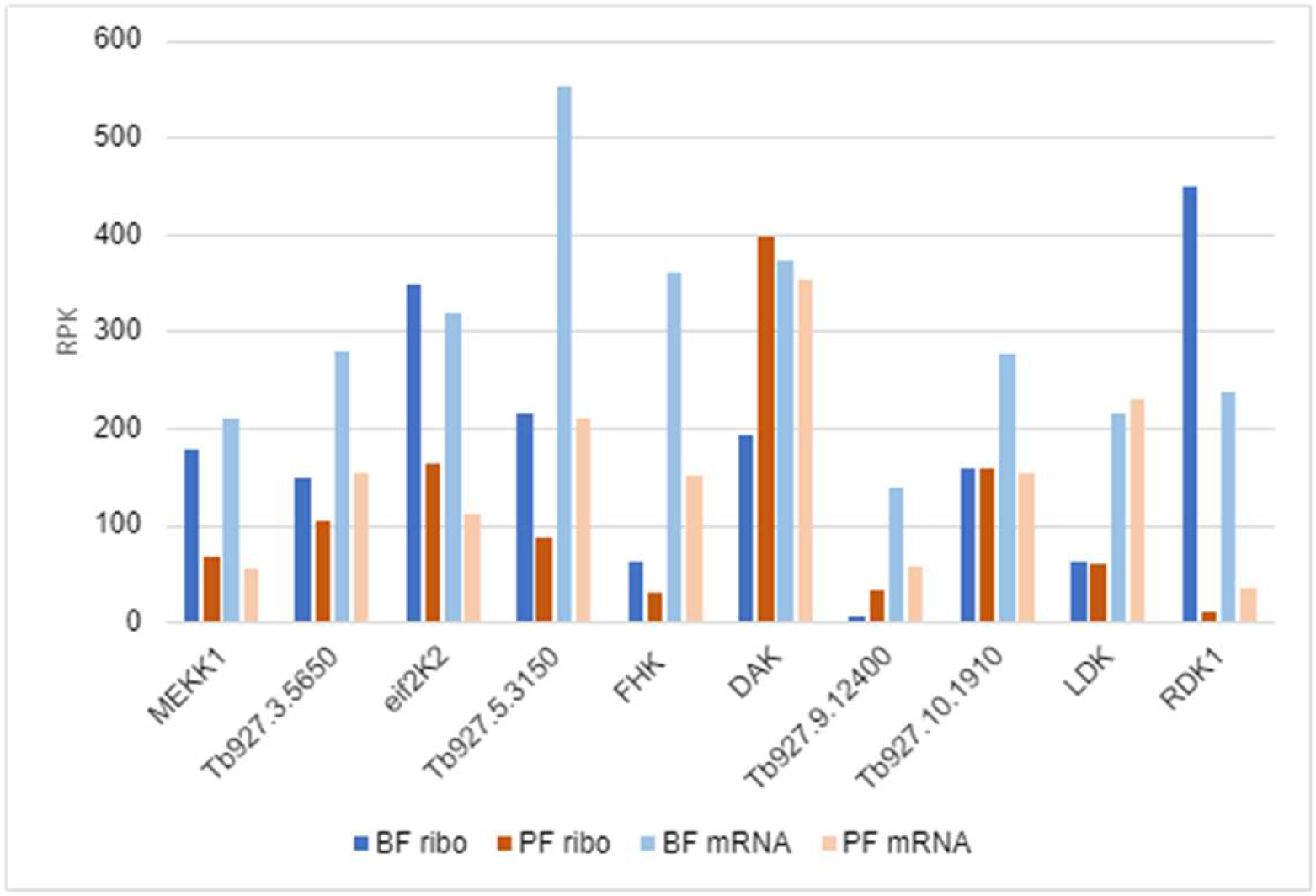
Protein production and mRNA levels of TMD PKs. Protein production was measured by ribosome profiling and mRNA levels were measured in the same biological samples (three replicates each). Data is shown as reads per kb and is derived from [25].

### Subcellular localization

Previous work localized *T. brucei* LDK to the lipid droplet membrane [10] and eIF2K2 to the flagellar pocket [11] (and its *T. cruzi* orthologue to endosomes [13]). Additionally, the protein encoded by Tb927.10.1910 was localized to the cytoplasm and endocytic organelles in PF in the TrypTag project [26]. We determined the localization of C-terminally tagged versions of the remaining transmembrane PKs in single marker strain BF. A western blot (Fig. S2) shows the migration of the tagged proteins in SDS gels, which corresponded reasonably well to the predicted sizes given in Table S1. In the absence of antibodies specific to individual PKs, it is not possible to predict whether the expression levels of the tagged protein are comparable to the endogenous levels. However, the very low levels of FHK and Tb927.9.12400 revealed by ribosome profiling compared to their ready detection by western blot (Fig. 3) suggest that the tagged versions of those PKs may be over-expressed.

#### Many tagged TMD PKs localize to the ER

As summarized in Table 1 and shown in Fig. 4, five of the tagged PKs with predicted TMDs colocalized with the ER marker BiP. None of these PKs is classified as related to the mammalian ER kinase PEK [7]. Staining of the tagged proteins is evident around the body of the cell but does not extend along the flagellum after it passes the anterior end of the cell body. As with BiP, some perinuclear staining is also seen; in most cases this is fainter than the peripheral staining. RDK1 was previously shown to be associated with membranes and localized to the periphery of the parasite [9]; our colocalization studies indicate that this peripheral localization is the ER. While it is clear that these PKs enter the secretory system, the ER may not be their final destination, which could be masked by potential over-expression.

**Table 1.** Summary of localization studies

| GeneID/name | TrypTag<br>PF<sup>a</sup>, endogenous | other studies | This study<br>BF, ectopic,<br>C-terminal tag |
| --- | --- | --- | --- |
| Tb927.2.2720<br>MEKK1 | N -faint<sup>b</sup>, cytoplasm and flagellum;<br>C - faint, cytoplasmic, reticular |  | Flagellum, flagellar pocket;<br>similar in PF |
| Tb927.3.5650 | nd |  | ER |
| Tb927.4.2500<br>eIF2K2 | N - very faint, cytoplasmic puncta;<br>C - nd | Flagellar pocket and<br>endosomes [11] | nd |
| Tb927.5.3150 | N- nd; C – reticulated with<br>occasional bright puncta |  | ER |
| Tb927.7.5220<br>FHK | nd |  | At or near flagellar pocket,<br>similar in PF |
| Tb927.9.3120 | nd |  | ER |
| Tb927.9.12400 | nd |  | ER |
| Tb927.10.1910 | N - faint, endocytic and cytoplasm;<br>C -ND |  | nd |
| Tb927.11.8940<br>LDK | nd | Lipid droplets [10] | nd |
| Tb927.11.14070<br>RDK1 | nd | Periphery [9] | ER |

<sup>a</sup> TrypTag: in situ tags the test protein with neon-green at the N- or C-terminus of the protein for expression in PF [26].
<sup>b</sup>Faint, 10-20<sup>th</sup> percentile of all lines expressing tagged proteins; very faint, <10<sup>th</sup> percentile. Note that these are all fainter than the parental line, but in cases where proteins are localized to specific organelles, localization can often be visualized.

**Figure 4.**
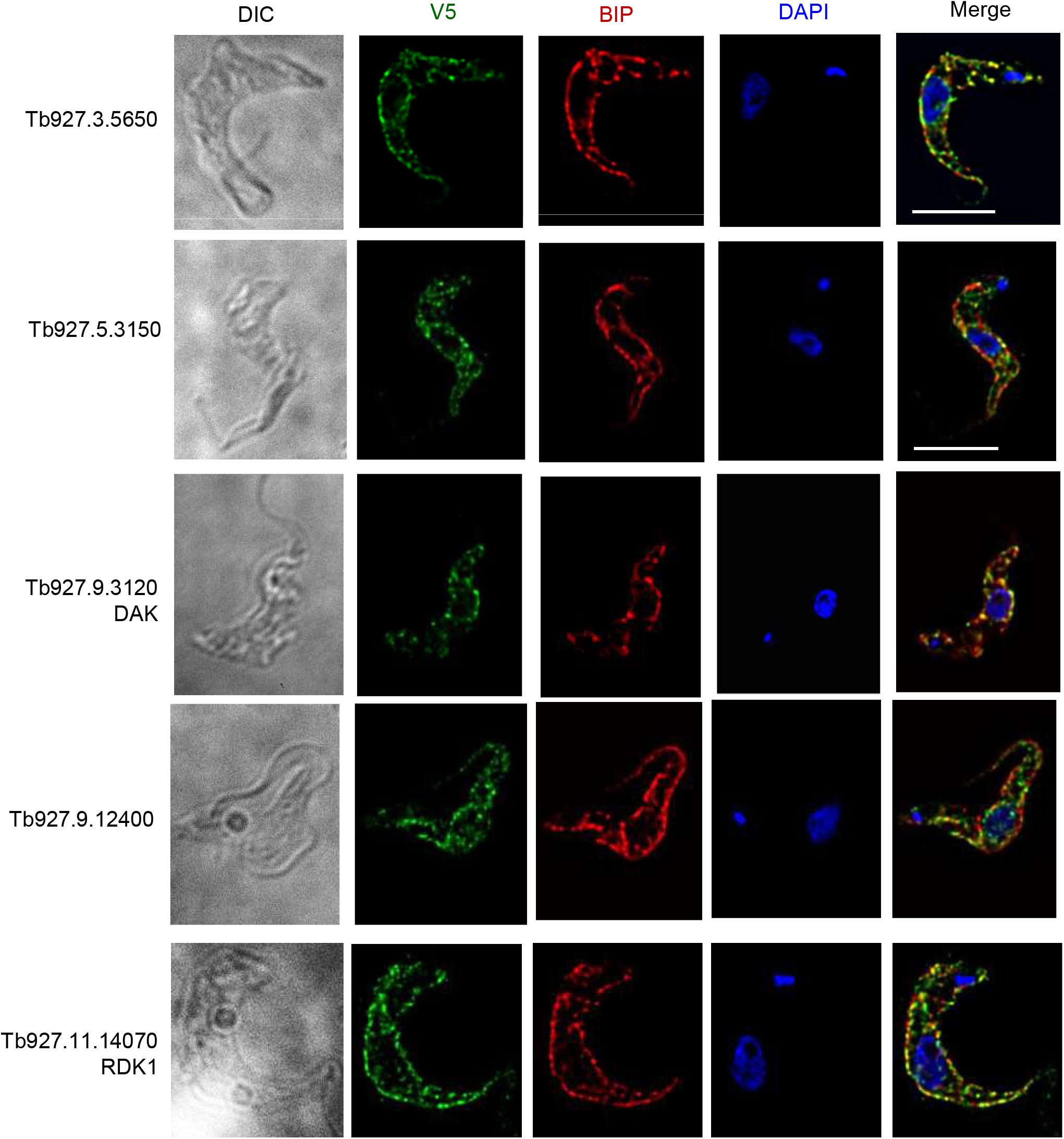
Localization of V5-tagged predicted membrane kinases to the ER in bloodstream form *T. brucei*. BF were fixed and stained with primary antibodies. Bound antibodies were revealed with secondary fluorochrome-coupled antibodies. Green, V5-tagged proteins, Red, anti-BiP, Blue, DAPI; Merge, all three colors. Each series shows a single deconvolved plane, except the DIC image, which is a projection. All images are shown at the same magnification. Bar, 5 μm.

#### MEKK1 is an integral membrane protein of the flagellum and flagellar pocket with kinase activity

Immunofluorescence analysis showed that MEKK1-V5 is present along the flagellum, where it aligns with PIFTC3, a component of the flagellar protein transport system [27] and is clearly distinct from the flagellar K+ channel that also extends along the flagellum [28]. We also observed staining adjacent to the kinetoplast, which was distinct from, but adjacent to, the depot of PIFTC3. When two kinetoplasts were seen, two puncta of both PIFTC3 and MEKK1 were visible. Immunoelectron microscopy was performed using PF expressing MEKK1 C-terminally tagged with HA epitopes. As shown in Fig. 6, gold particle staining was present at the flagellar pocket (A, B) but not at the kinetoplast (C). In a dividing cell, both flagellar pockets were stained. MEKK1-HA was also seen along the flagellum (Fig 6E). The MEKK1-HA at the flagellar pocket may represent molecules in transit to the flagellum or a final destination, as is seen for PIFTC3 [27] and (Figure 5).

**Figure 5.**
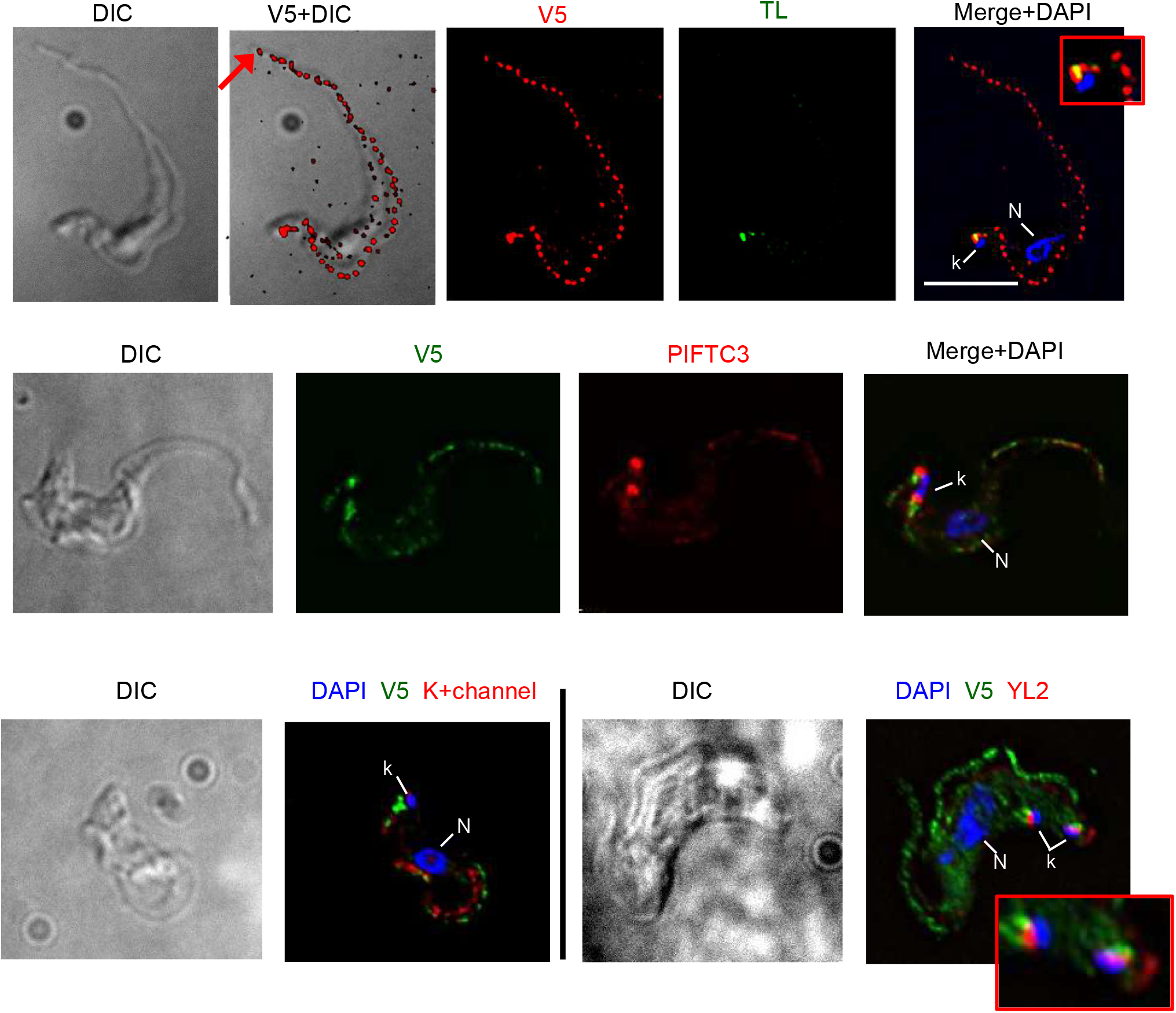
Immunofluorescence localization of MEKK1-V5 (Tb927.2.2720) in *T. brucei* bloodstream forms. <u>Top</u>, co-staining with tomato lectin (TL, green) at 4°, which marks the flagellar pocket. Note that punctate staining continues to the end of the flagellum. <u>Middle</u>, co-staining with antibody to T. brucei PIFCT3 (red), a component of the flagellar transport system<u>. Bottom</u>, co-staining with antibody to a K+ channel protein of T. cruzi (left, red) and with monoclonal antibody YL2 (right, red) which detects basal bodies. The latter shows a cell with two kinetoplasts and appears to be in early mitosis. Each series shows a single deconvolved plane, except DIC images, which are projection. All images, except enlargements, are shown at the same magnification (bar, 5 μm). Enlargements show detail near the kDNA. N, nucleus; k, kDNA.

**Figure 6.**
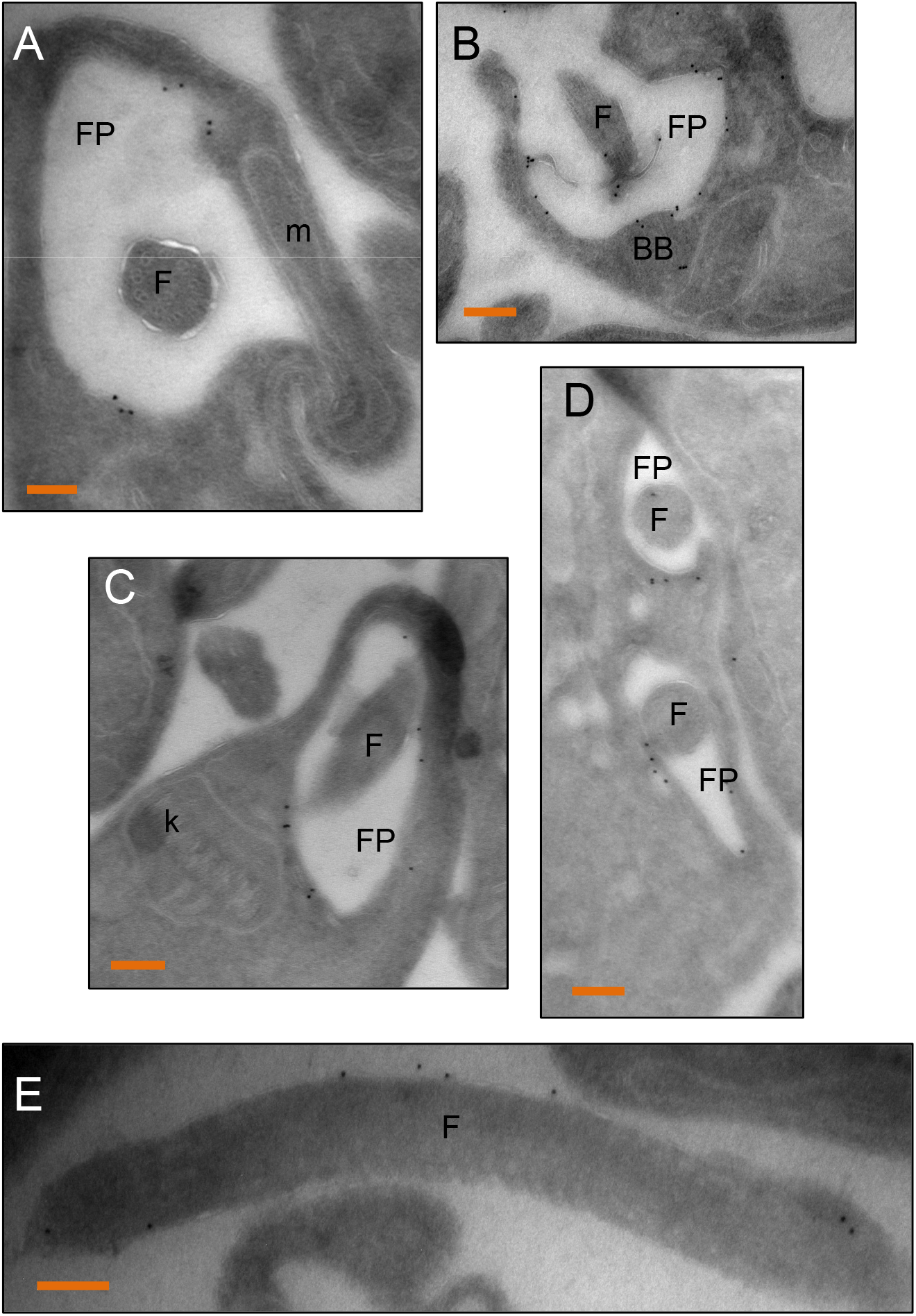
Localization of MEKK1-HA to the flagellum and flagellar pocket by immunoelectron microscopy. PF expressing MEKK1 C-terminally tagged with HA epitopes were stained with anti-HA, and staining revealed by protein-A gold 10 nm particles. A) The gold staining is on the flagellar pocket (FP) membrane, flagellum (F), but absent from the mitochondrion (m). B) staining is observed at the base of the flagellar pocket at the basal bodies (BB). C) no staining of the kinetoplast (k) is seen. D) in a dividing cell, both flagellar pockets are stained. E) the external portion of the flagellum is also stained. Bars, 100 nm.

To assess membrane association, we conducted cell extraction and fractionation of PF expressing MEKK1-HA. The full-length protein migrated anomalously, somewhat faster than the 220 kDa marker (the tagged protein is predicted to be 167 kDa). Membrane proteins are known to exhibit anomalous migration on SDS-PAGE gels, migrating either slower or faster than predicted, with apparent molecular masses averaging 80-113% of the predicted value [29]. The ∼95 kDa V5-tagged fragment can be disregarded since it is not seen in cells lysed directly in SDS sample buffer, and therefore represents a degradation fragment. As shown in the immunoblot in Fig. 7, upon treatment of PF with digitonin to release the cytosol, the intact kinase remained in the pellet fraction (P1). Subsequent treatment of the digitonin pellet with sodium carbonate, pH 11, showed MEKK-V5 in the pellet (P2), demonstrating that it is an integral membrane protein. Marker proteins PGKB (cytosolic), PGKA (glycosomal which shows partial membrane association) [30] and vacuolar pyrophosphatase, TcVSP) (integral vacuolar membrane) [31] verified the fractionation procedure.

**Figure 7.**
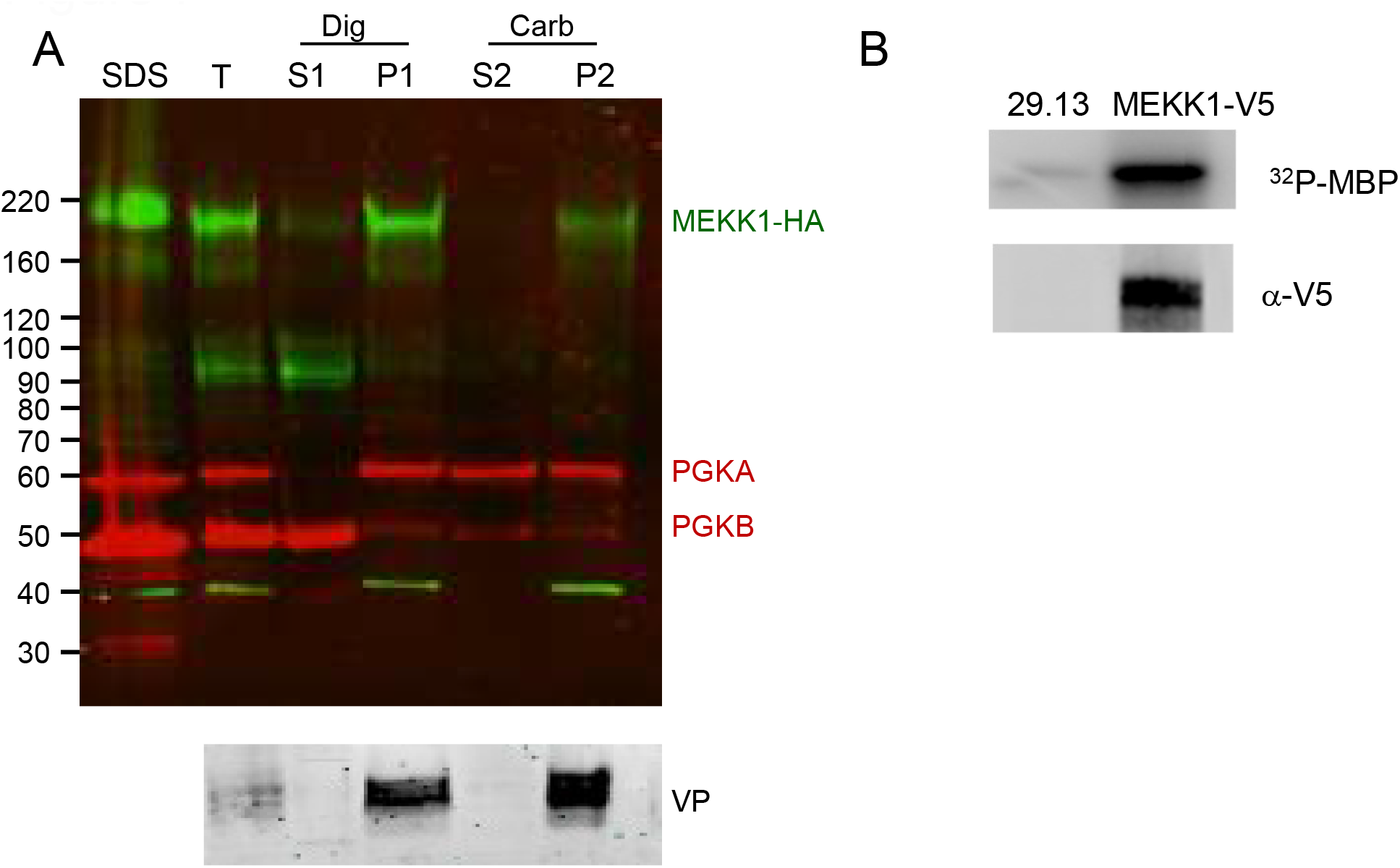
MEKK1 is an integral membrane protein which has protein kinase activity. A) Cell fractionation. PF cells expressing MEKK1-HA were fractionated by digitonin treatment followed by carbonate extraction. After separation by SDS-PAGE and blotting, the blots were probed with mouse anti-HA (green) and rabbit anti-phosphoglycerate kinase (red) which detects the 56 kDa glycosomal form (PGKA) and the 47 kDa cytosolic isoform (PGKB). A parallel blot was also incubated with antibody to the acidocalcisome membrane protein vacuolar phosphatase (VP). SDS, SDS lysate; T, total after incubation with digitonin; S1, digitonin supernatant; P1, digitonin pellet; S2, carbonate supernatant; P2, carbonate pellet. Lanes contain fractions from 3×106cells. B) Kinase activity. Immunoprecipitated MEKK1 from PF was incubated with γ 32P-ATP and the exogenous substrate myelin basic protein (MBP), followed by SDS-PAGE and transfer to nitrocellulose. A control immunoprecipitation used the untransfected parental line (29.13). <u>Top:</u> phosphorimaging demonstrates phosphorylation of MBP. <u>Bottom:</u> the blot was probed with anti-V5 mAb demonstrating the pulldown of the tagged protein.

To test for catalytic activity, anti-V5 immunoprecipitates from PF expressing MEKK1-V5, as well as from the untransfected parental 29.13 line, were incubated with the exogenous substrate myelin basic protein (MBP) in the presence of γ32P-ATP. The proteins were then separated by SDS-PAGE and analyzed by phosphorimaging (Fig. 7B, top) and western blot (Fig. 7B, bottom). MBP was phosphorylated in the sample containing MEKK1-V5, but not the untransfected control, thereby demonstrating that MEKK1 can phosphorylate an exogenous substrate.

#### FHK is an integral membrane protein of the flagellar pocket with kinase activity

Like MEKK1, FHK-V5 demonstrated staining near the kinetoplast, close to PIFTC3 upon immunofluorescence analysis of BF (Fig. 8A). However, staining did not extend along the flagellum and little co-localization with BiP was observed (Fig. 8B). Live PF expressing MEKK1-HA were incubated with biotinylated tomato lectin at 4° C and then fixed for anti-HA staining (Fig. 8C). The low temperature allows lectin binding to the flagellar pocket without internalization into the parasite. The HA staining was adjacent to and possibly overlapping the tomato lectin signal. Fig. 8D depicts a PF parasite stained for DNA and FHK. Individual deconvolved planes showed FHK between the nucleus and kinetoplast in what appears to be a basket-like structure.

**Figure 8.**
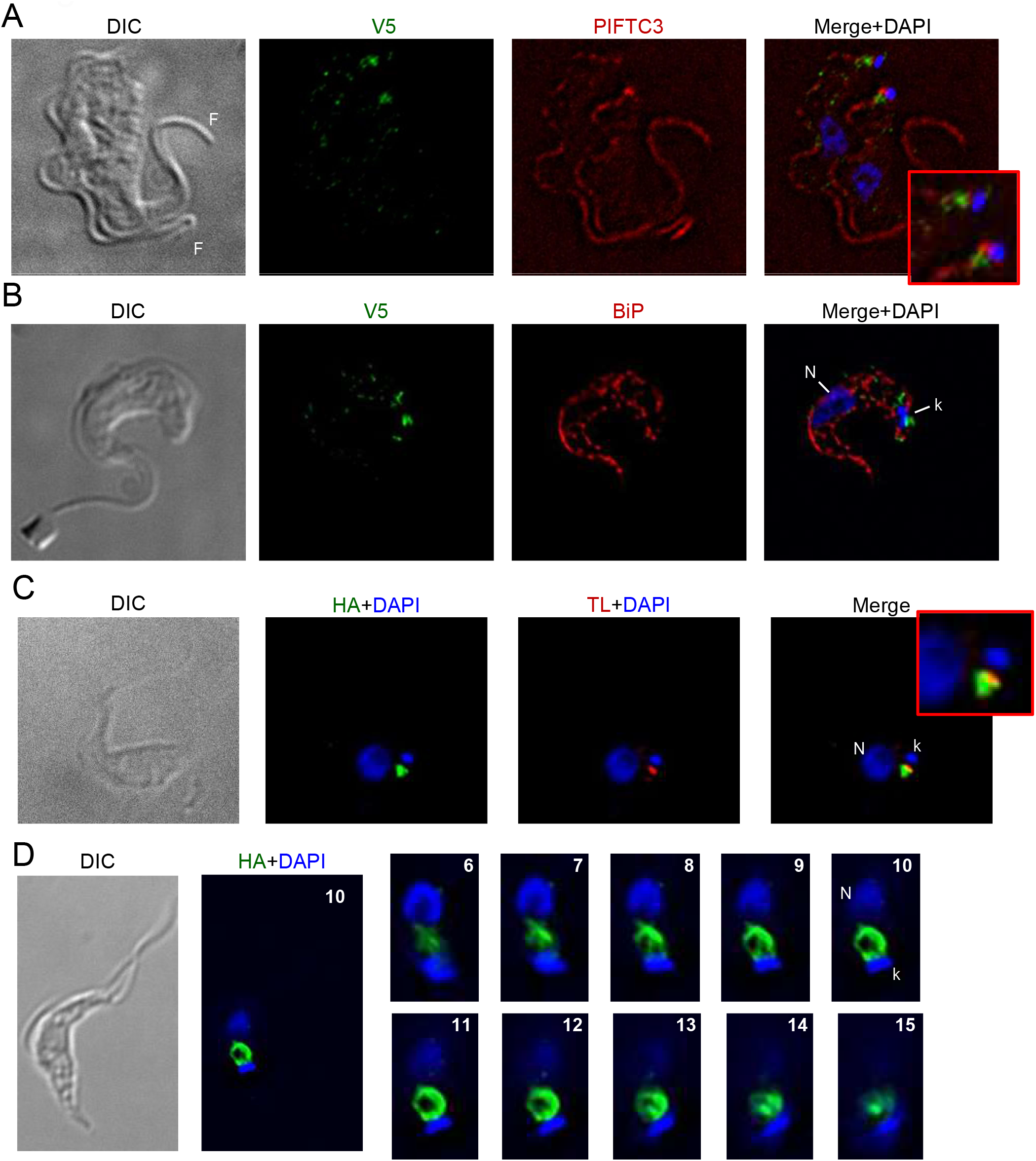
Predominant localization of epitope-tagged FHK near the kinetoplast. **A, B)** BF expressing FHK-V5 and co-stained with DAPI, anti-V5 antibodies, and either anti-PIFTC3 (A) or anti-BiP (B). Enlargement shows region around kDNA. The cell in (A) has duplicated both its nucleus (N) and kDNA (k) and is undergoing cytokinesis as evidenced by the presence of two flagella (F). The cell in panel B has an elongated, slightly V-shaped kinetoplast, indicating it is undergoing kinetoplast division and is in nuclear S phase. Note that a point of FHK-staining occurs at either pole of the elongated kDNA. **C)** PF expressing FHK-HA were incubated with biotinylated-tomato lectin (TL) at 4o and fixed prior to staining with DAPI, avidin and anti-HA. Enlargement shows region around kDNA. **D)** Whole cell and enlarged serial planes (numbered) of a DAPI-stained PF cell expressing FHK-HA. All images (except enlargements) are at the same magnification. Bar = 5 μm.

Immunoelectron microscopy of PF demonstrated that the protein was localized to the flagellar pocket but not to the flagellum or kinetoplast (Fig. 9A, enlarged in 9a), and occasionally staining is seen more prominently on one side of the flagellar pocket (Fig. 9B, enlarged in 9b). Sometimes staining was observed on vesicles close to the flagellar pocket, but only sporadically outside the region of the pocket (Fig. 9 C, D).

**Fig. 9.**
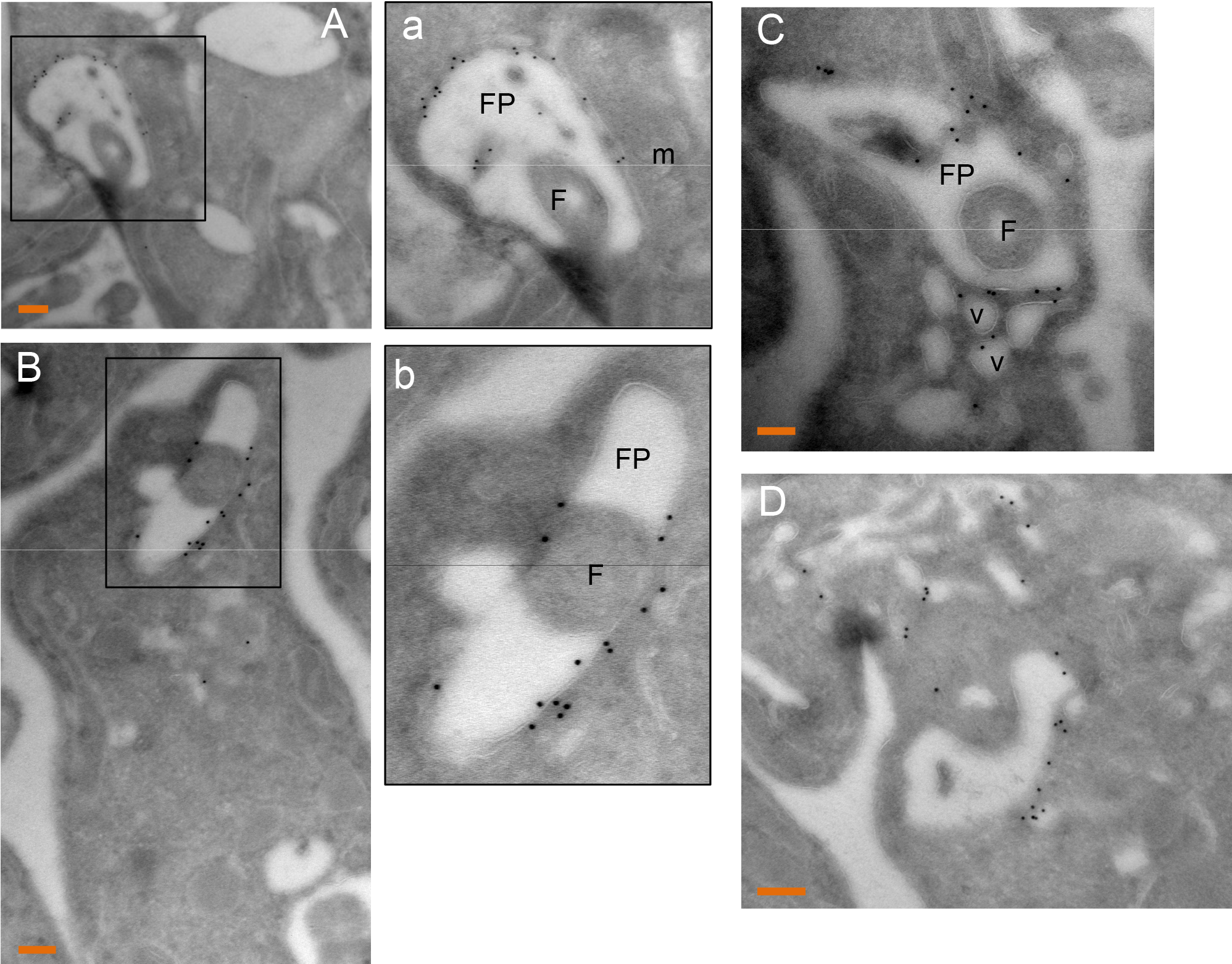
Localization of FHK-HA to the flagellar pocket by immunoelectron microscopy. Procyclic form parasites expressing FHK C-terminally tagged with HA epitopes were stained with anti-HA revealed by protein-A gold 10 nm particles. A) The gold staining is on the flagellar pocket (FP) membrane but absent from the flagellum (F) and mitochondrion (m). Enlargement of the marked region is shown in panel a. B) Staining is predominantly at the flagellar pocket, and often one side shows more abundant FHK. Enlargement of the marked region is shown in panel b. C) The staining is sometimes seen on vesicles close to the FP. D) Occasional parasites showed staining outside the flagellar pocket. Bars, 100 nm.

PF expressing FHK-HA were fractionated as described above for MEKK1. Full-length FHK-HA was in the digitonin pellet, and the carbonate-insoluble fraction, indicating it is an integral membrane protein (Fig. 10A). As with MEKK1, the smaller degradation fragment can be disregarded since it was absent from cells lysed in SDS sample buffer. To test for catalytic activity, immunoprecipitates from PF expressing FHK-HA, as well as the untransfected, parental single marker line, were assayed as described for MEKK1. As shown by phosphorimaging (Fig. 10B, top) and western blot (Fig. 10B, bottom), MBP was phosphorylated in the sample containing FHK-HA, but not the untransfected control, providing evidence of FHK catalytic activity.

**Figure 10.**
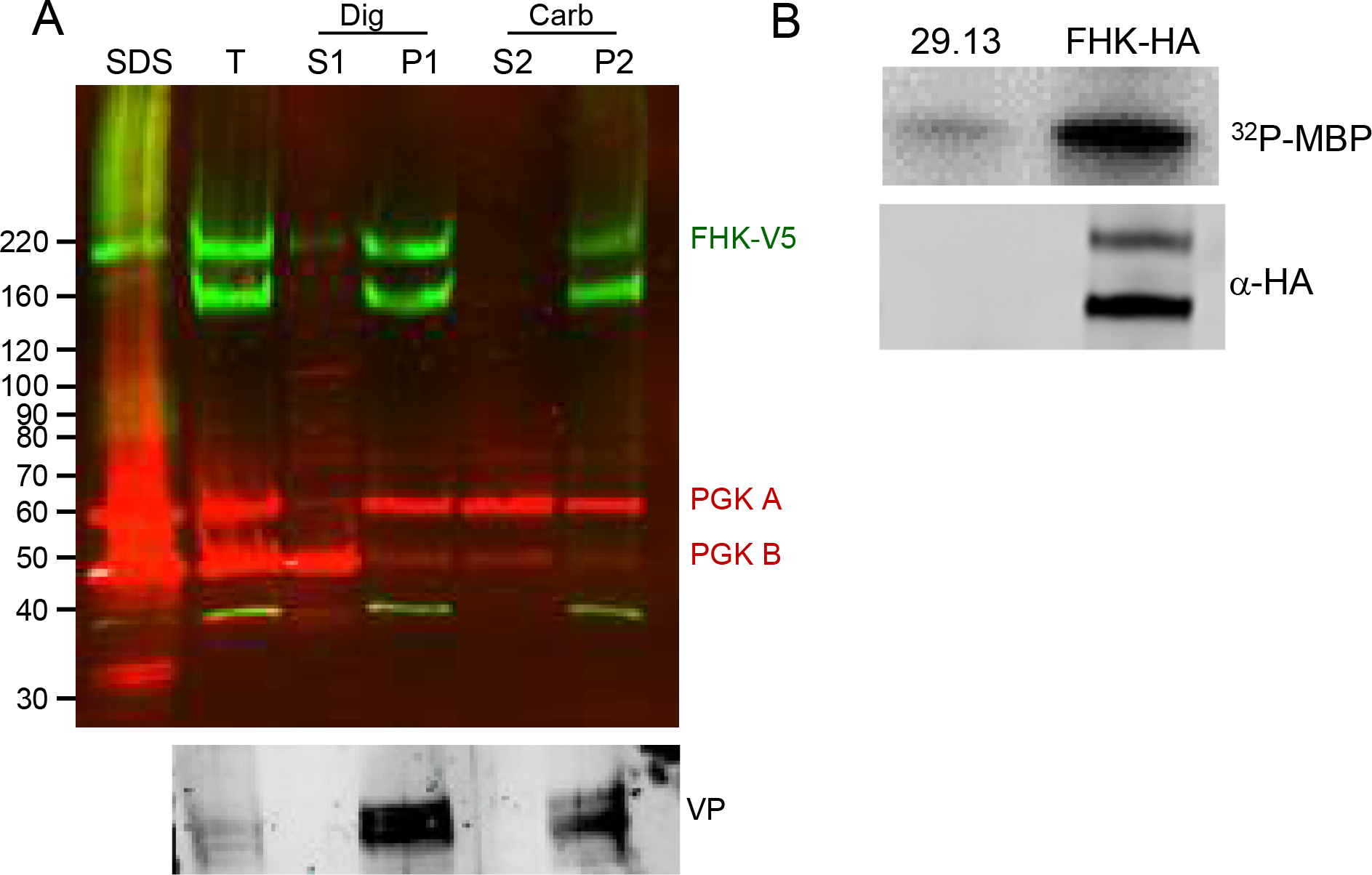
FHK is an integral membrane protein with protein kinase activity. **A)** Cell fractionation and immunoblotting were as described in Fig. 7. **B)** Kinase activity as described in Fig. 9. FHK-HA was immunoprecipitated from PF and incubated with γ 32P-ATP and MBP. A negative control immunoprecipitation was performed using the untransfected parental line 29.13. Top: phosphorimaging demonstrates phosphorylation of MBP. Bottom: the blot was probed with anti-HA mAb demonstrating the pulldown of the tagged protein.

## Discussion

### A plethora of transmembrane domains

In contrast to metazoan kinomes, which have large numbers of TMD PKs, protozoan kinomes vary widely in their representation of TMD PKs. At one extreme, the *Giardia lamblia* genome encodes only two predicted transmembrane PKs (two other PKs have predicted TMDs unlikely to be authentic since they punctuate the catalytic domain). At the other extreme is *Entamoeba histolytica* with more than 80 predicted TMD PKs, most of which are related at the sequence level and some of which are likely non-catalytic [32]. Like *E. histolytica, Plasmodium falciparum* has evolved a unique set of PKs, the FIKK kinases, that comprise the majority of the parasite’s 17 PKs with predicted TMDs. The FIKK kinases bear the signature sequence for export out of the parasite [33], and at least some of them are important in remodeling the host erythrocyte [34, 35]. Like *P. falciparum*, trypanosomatids fall between extremes of *G. lamblia* and *E. histolytica*, with a paucity of PKs that bear TMDs compared to metazoans. These trypanosomatid PKs are classified firmly with serine/threonine kinases as opposed to tyrosine kinases characteristic of transmembrane PKs of animals [7]. Just as interesting, most of these trypanosomatid PKs are predicted to have multiple membrane spanning domains, a feature that is extremely rare in eukaryotes. To our knowledge, a single ePK domain protein in eukaryotes has been biochemically demonstrated to have more than one TMD, lemur tail kinase 2 [36], but other members of that family of PKs are also predicted to have two TMDs [37]. Interestingly, in perusing the *E. histolytica* genome database, we noted a few PKs with two likely TMDs. Another interesting observation is that many bacterial histidine kinases with several TMDs have been identified in genome-wide bioinformatic analyses [38]. These include the 5-TMD bearing *Staphylococcus aureus* LytS histidine kinase which acts as a sensor of membrane potential [39]. Another example is *Bacillus subtilis* DesK, in which TMDs are involved in signaling temperature-induced changes in membrane fluidity [40]. Thus, TMDs can modulate kinase activity in response to environmental changes even when no extracellular ligand binding domains are present. It will be interesting to determine if any of the TMDS in the *T. brucei* PKs examined here have such effects.

### Flagellar and flagellar pocket localization and function

The localization of MEKK1 and FHK to the flagellum and flagellar pocket respectively raises the question of what biochemical processes they may regulate. Initiation of stumpy development appears to be triggered by the uptake of oligopeptides by the transporter GPR89 [41]. MEKK1 acts early in the developmental pathway, which involves multiple additional kinases [8]. How the oligopeptide signal intersects with MEKK1 is not clear given that GPR89 is distributed across the plasma membrane and is not specifically localized to, or perhaps not present at all, at the flagellar membrane. However, other molecules that may be involved in development are also localized to the flagella, such as adenylate cyclases and phosphodiesterases that are present on the flagellum, as well as a calcium channel [42].

In trypanosomatids, the flagellar pocket is the site of the essential processes of exocytosis and endocytosis. Therefore, the localization of FHK to the flagellar pocket could suggest a function in these processes. RNAi data suggests that FHK is dispensable in BF [9, 43] (also corroborated by our preliminary data), so it is unlikely that it functions as a global regulator of such essential processes. Instead, its function may be focused on specific conditions (e.g., stress response) or certain pathways within the broader categories of exocytosis, endocytosis and recycling of membrane molecules. Because only the flagellar pocket is free of the densely packed, GPI-anchored variant surface protein coat which covers the surface of the parasite and bears many distinct proteins, FHK could also be positioned at the flagellar pocket to sense interaction with transmembrane proteins or even to biophysical changes in this distinct membrane domain, such as those sensed by LytS and DesK.

### Signaling domains of likely bacterial origin

There are numerous genes in trypanosomatids, especially those encoding metabolic enzymes, which are more closely related at the amino acid sequence level to bacterial than to eukaryotic homologues, providing evidence of lateral gene transfer. Our HHpred analysis showed that two *T. brucei* PKs have regions predicted at >95% probability to fold similarly to bacterial signaling modules. FHK has a region resembling ligand-sensing domains found on chemotaxis proteins and certain histidine kinases, but not on eukaryotic proteins. DAK has cassette bearing four regions common to hybrid histidine kinases: a sensor, a dimerization/phosphoacceptor, a catalytic domain and a receiver domain. These modular structures are conserved in genomes across trypanosomatids and in the bodonid *Bodo saltans*, ruling out an artifactual explanation and arguing for a functional and possibly regulatory role. While this combination of domains is unusual, Uniprot searches do identify predicted proteins that possess both an ePK domain and a histidine kinase cassette. Most of these predicted proteins are bacterial, but some are found in eukaryotes (predominantly fungi). Those in eukaryotes almost exclusively have hybrid type histidine kinase domains (i.e, with a receiver domain on the same molecule) as is seen on DAK. Few (if any) of these histidine-ePK kinases have been explored functionally to understand how the domains interact and influence the signaling properties of the protein. DAK’s potential for histidine kinase activity that phosphorylates the DHp region appears low since the conserved histidine is mutated. However, the aspartic acid in the receiver domain that accepts the phosphate is still present. Clearly additional work will need to be done to understand the roles of the two ancestral kinase regions on the kinase activity and ultimately function of DAK [25].

## Materials and Methods

### Parasites

The work described uses PF 29-13 and single marker BF lines, which are derivatives of the *T. brucei* 427 strain [44]. Both lines express T7 RNA polymerase and the tetracycline (Tet) repressor, allowing for Tet-regulated expression of transfected genes. PF were grown in SDM-79 (JRH Biosciences) supplemented with 15% fetal calf serum containing 15 μg/ml G418 and 50 μg/ml hygromycin to maintain the T7 RNA polymerase and Tet repressor genes. BF were grown in HMI-9 with 2.5 μg/ml G418. Plasmids were transfected into PF as described [44] and transfectants were selected with 1 μg/ml puromycin. BF were transfected as described [45] and modified by [46]. BF transfectants were selected with either 5 μg/ml hygromycin or blastocidin S depending on the plasmid vector.

### Plasmids and Cloning

For expression of V5-tagged proteins in BF, we generated the plasmid pT7-3V5-Hyg by replacing the GFP and lacZ stuffer region of pT7-GFP (kindly supplied by David Horn) [47] with the multicloning site and 3V5 epitopes derived from pLEW79-3V5-Pac [10]. Expression was driven by a tetracycline (Tet) regulated T7 promoter and the 3’ UTR was derived from aldolase. For expression of HA-tagged proteins in PF, either pLEW79-3V5-PAC or plasmid pLEW100v5-3HA-BSD were utilized. The latter was created by replacing the luciferase gene in pLEW100v5-BSD (kindly supplied by George Cross) with the multicloning sites derived from pLEW79-3V5-PAC and three HA epitope tags. Expression was driven by a tetracycline (Tet) regulated PARP promoter and the 3’ UTR was derived from aldolase. The coding sequences for the PKs were amplified and PCR products were digested with the restriction enzymes noted in the primer and ligated into appropriately digested vectors. The expression construct for Tb927.9.3120 also included the endogenous splice acceptor. All primers are described in Table S2.

### Western blots

Cell lysates were generated, and proteins resolved by SDS-PAGE and transferred to nitrocellulose as described [48]. V5-tagged proteins were detected with mouse monoclonal anti-V5 antibody (ThermoFisher) at 0.3 μg/ml. HA-tagged proteins were detected with mouse (Covance) or rat (Roche) monoclonal anti-V5 antibodies at 1 μg/ml or 12.5 ng/ml respectively. Bound antibodies were revealed with IRDye 800CW dye-conjugated goat anti-mouse Ig (Li-COR) at 25 ng/ml and data was visualized on a Li-COR Odyssey.

### Immunofluorescence analysis

IFAs were performed as previously described [48]. In brief, BF parasites were prefixed in 4% paraformaldehyde for 5 min on ice prior to placing on poly-L-lysine coverslips. The V5 epitope tag was detected by using mouse monoclonal anti-V5 antibody (Invitrogen) at 1 ug/ml, followed by goat anti-mouse IgG conjugated with fluorescein isothiocyanate (FITC) (Southern Biotechnology). Additional antibodies used for colocalization included antibodies directed against BiP [49] (kindly supplied by Jay Bangs), used at 1:400; PIFTC3 [27] (kindly supplied by Elisabetta Ullu), used at 1:1000; TcCat [28] (kindly supplied by Roberto DeCampo), used at 1:250; YL1/2 (Gene-Tex), used at 1 μg/ml. To visualize the flagellar pocket, pre-cooled cells were incubated at 4°C with biotinylated tomato lectin (Vector Laboratories) at 2 μg/ml. After fixation, tomato lectin was visualized with 2 μg/ml streptavidin conjugated to either Texas Red or Alexa Fluor 488 (ThermoFisher). Dye-conjugated secondary antibodies (Southern Biotechnology) were used at 2 μg/ml. Images were scaled to span the graph of signal intensity of the image field.

### Immunoelectron microscopy

Preparations of parasites were fixed in 4% paraformaldehyde (Electron Microscopy Sciences, PA) in 0.25 M HEPES (pH 7.4) for 1 h at room temperature, then in 8% paraformaldehyde in the same buffer overnight at 4oC. They were infiltrated, frozen and sectioned as previously described [50]. The sections were immunolabeled with rat anti-HA antibodies (1:50 in PBS/1% fish skin gelatin), then with anti-mouse IgG antibodies, followed directly by 10 nm protein A-gold particles (Department of Cell Biology, Medical School, Utrecht University, the Netherlands) before examination with a Philips CM120 Electron Microscope (Eindhoven, the Netherlands) under 80 kV.

### Cell fractionation and Kinase assays

PF expressing HA-tagged proteins washed in PBS containing 10 mM glucose. They were permeabilized with digitonin and the organellar pellet was extracted with sodium carbonate as described [51] except that Complete Protease tablets (Roche) were included for both digitonin permeabilization and carbonate extraction, to prevent protein degradation.

Cell lysates from PF cells expressing either V5-tagged MEKK1 or HA-tagged FHK along with the untransfected parental line 29.13 were prepared as described [52]. Epitope tagged proteins were immunoprecipitated from cell lysate prepared from 10^8^ cells. Kinase assays were done as described [53] using ^32^PγATP with myelin basic protein as an exogenous substrate. Reactions were resolved by SDS-PAGE and transferred to nitrocellulose and labeled proteins detected by phosphorimaging. Blots were subsequently probed with antibodies directed against either V5 or HA.

## Acknowledgements

This work was supported in part by a grant 5R21AI101424 from the National Institutes of Health and by Seattle Children’s Research Institute.

## Author contributions

Concept and experimental design were done by B.C.J and M.P. Experiments were performed by B.C.J, P.V. and J.F except electron microscopy done by I.C. Manuscript was written by M.P. and B.C.J. All authors have reviewed the manuscript.

## List of supplementary information

### Supplementary tables

Table S1. TriTryp orthologues - transmembrane domains, signal sequences, and additional domains

Table S2. Primers

### Supplementary figures

Fig. S1. RDK1 structural similarities identified by HHpred

Figure S2. Western blot showing expression of epitope-tagged PKs in *T. brucei* bloodstream forms.

