## Supplementary Tables for "Unusual Features of the Membrane Kinome of *Trypanosoma brucei*"

| Table S1. TriTryp orthologues - transmembrane domains (TMHMM predicted), signal sequences, and additional domains | | | | | | | |
|---|---|---|---|---|---|---|---|
| *T. brucei* | MW kDa | TMD and SS* | *T. cruzi** | TMD and SS** | *L. major* | TMD and SS** | additional domains/comments |
| Tb927.2.2720 (MEKK1) | 162 | 607-629, 634-656 | TcCLB.508739.70 | 757-779, 788-810 | LmjF.02.0570 | 578-600, 607-629 | |
| Tb927.3.5650* | 104 | 47-69,103-125 | TcCLB.510645.9 | 30-52,82-104 | LmjF.21.0130 | 44-66, 99-121 | nucloeotide cyclase domain |
| Tb927.4.2500 (eIF2K2) | 112 | 1-26 (ss**), 4-23, 465-487 | TcCLB.505559.129 | 1-24 (ss), 7-29 | Lmjf.34.2150 | none annotated | CCTOP predicts TMDs in L. major |
| Tb927.5.3150 | 178 | 1-22 (ss), 7-29, 442-464, 607-629 | TcCLB.503955.100 | 1-22 (ss), no TMD | LmjF.08.0530 | 1-21 (ss), 85-101. | CCTOP predicts TMDs on T. cruzi orthologue. Kelch domains on T. brucei and T. cruzi proteins. L. major ortholog lacks the N-terminal ~370 aa, as well as the Kelch domains |
| Tb927.7.5220 (FHK) | 182 | 20-42, 57-79, 141-163, 200-222, 235-257 | TcCLB.506825.180 | 13-35, 50-72, 204-226, 236-258, 385-407 | LmjF.06.0640 | 199-221, 225-247, 321-343, 395-417, 424-446 | Forkhead domain, HHpred similarities to PAS |
| Tb927.9.3120 | 112 | 146-161, 181-203 | TcCLB.510565.70 | 150-172, 185-207 | LmjF.26.1730 | 153-175,182-204 | conserved PAS domains; HHpred similarities to histidine kinase DHp, catalytic and receiver domains |
| Tb927.9.12400 | 148 | 263-285, 323-345 | TcCLB.510741.70 | 266-288, 331-353 | none | N/A | nucleotide cyclase domain |
| Tb927.10.1910 | 168 | 218-240 | TcCLB.510121.150 | 1-34 (ss), 174-196, 499-521 | Lm.21.0130 | 315-337, 647-669 | CCTOP predicts TMD on T. cruzi orthologue. nucleotide cyclase domain |
| Tb927.11.8940 (LDK) | | 433-455 | TcCLB.511801.14 | none | LmjF.28.2000 | 533-555 | membrane monolayer |
| Tb927.11.14070 (RDK1) | 131 | 88-107, 111-128, 452-474 | TcCLB.511727.210 | 92-114, 453-475 | LmjF33.2290 | 380-403,785-808 | HHpred similarities to sensor/receptor domains and to adenylyl/gualyl cyclase domains. |

* Annotated on TriTrypDB via TMHMM and SP-HMM and SP-NN. T. brucei further analyzed with SignalP.

Table S2- Primers

| Primer | Sequence (5'-3') |
|---|---|
| MEKK1-AvrII-ATG | CCGCCTAGGATGCCCTTCGCGGCAAATGTTG |
| MEKK1-HindII-P4503M | CGGAAGCTTAACAACTCCCTGATCACCAAATG |
| Tb927.3.5650-AvrII-ATG | CCTAGGATGATAGAATATGCGTGTGGTTG |
| Tb927.3.5650-BamHI-P2787M | GGATCCCAAATGAAATTCGCTTCTCGAGATATTCCCAC |
| Tb927.5.3150-AvrII-ATG | CCTAGGATGGTTGTGCGGCCATTGTTCTG |
| Tb927.5.3150-BamHI-P4962M | GGATCCAAGCTCCTGGACATTACTGAAGAAG |
| FHK-AvrII-ATG | CGGCCTAGGATGACACAGCCCTCGGCAGATG |
| FHK-XhoI-P5034M | CGGCTCGAGATTACCTTCCACAGCAGCGTTAC |
| Tb927.9.12400-AvrII-ATG | CCTAGGATGTGTGCCTTAAGGGATATCGCGTCAAC |
| Tb927.9.12400-BglII-P4035M | AGATCTGTTTACCATAAAGTCATGGAACAAAAGCTCAG |
| DAK-AvrII-ATG | CCGCCTAGGATGGGACCAACATGTGATCGAATAG |
| DAK-BamHI-P3039M | CGCGGATCCCGAATGGGGTTGTGTGGTGATGGTAAG |
| RDK1-AvrII-ATG | CCGCCTAGGATGACGAAAGAGGATCACGGTG |
| RDK1-BamHI-P3585M | CGCGGATCCAACAAAAATGCATATGAAAGTAAACTGGATG |

Underlined bases are those used for inserting the gene into pT7-3V5-Hyg. Primer names include the restriction sites used for cloning. Bases preceding the restriction site were added to facilitate restriction digest of the PCR fragments.
