## Supplementary Figures for "Unusual Features of the Membrane Kinome of *Trypanosoma brucei*"

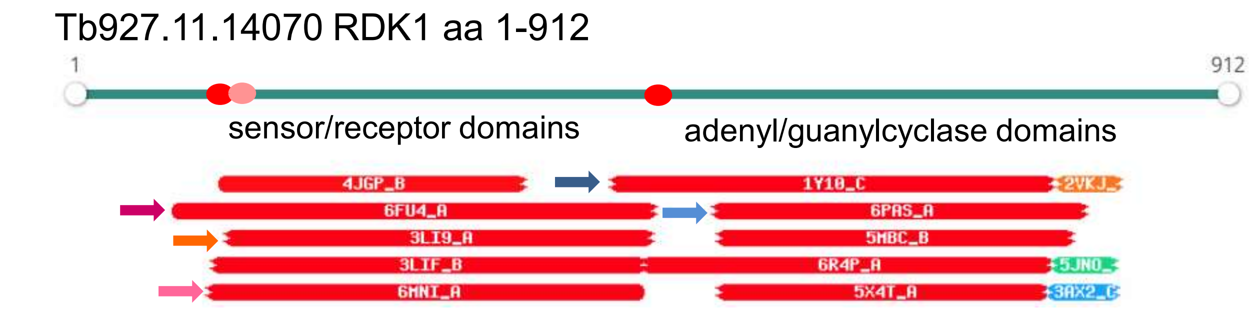


| Nr | Hit | Name | Probability | E-value | SS | Cols | Target Length |
|---|---|---|---|---|---|---|---|
| 1 | 1Y10_C | Hypothetical protein Rv1264/MT1302 (E.C.4.6.1.1); adenylyl cyclase fold, LYASE; HET: 1PE; 2.3A {Mycobacterium tuberculosis}; Related PDB entries: 1Y10_B 1Y10_D 1Y10_A 1Y11_A | 99.38 | 5e-11 | 23.9 | 277 | 407 |
| 2 | 6FU4_A | Chemotaxis protein; chemotaxis, Pseudomonas aeruginosa, signaling protein; HET: GOL, HSM, PO4, MSE; 2.45A {Pseudomonas aeruginosa PAO1}; Related PDB entries: 6FU4_B 6FU4_D 6FU4_C | 99.28 | 1.8e-9 | 27.6 | 319 | 345 |
| 3 | 6PAS_A | Soluble guanylyl cyclase alpha-1 subunit; Nitric oxide, cyclase, H-NOX, SIGNALING; HET: HEM; 5.1A {Manduca sexta}; Related PDB entries: 6PAT_A | 99.27 | 2.6e-10 | 22.9 | 243 | 699 |
| 4 | 3LI9_A | Hypothetical sensory transduction histidine kinase; PDC fold, SIGNALING PROTEIN; HET: MSE, BTB; 1.7A {Methanosarcina mazei} SCOP: d.110.6.4; Related PDB entries: 3LIB_B 3LIB_E 3LIB_J 3LIB_H 3LIB_F 3LIB_D 3LIB_G 3LIB_C 3LIB_I 3LIB_A 3LI8_A 3LIA_B 3LIA_A | 99.23 | 5.5e-9 | 27.1 | 276 | 291 |
| 5 | 3LIF_B | Putative diguanylate cyclase (GGDEF) with; PDC fold, SIGNALING PROTEIN; HET: MSE, MPD, CIT; 2.7A {Rhodopseudomonas palustris}; Related PDB entries: 3LIF_A | 99.2 | 1.8e-8 | 29.2 | 241 | 254 |

**Fig. S1. RDK1 shows evidence of additional signaling domains.** The C-terminal PK domain is not shown. Red ovals – transmembrane domains predicted by TMHMM and CCTOP. Each bar below the gene model represents the top hits by HHpred. The color of the arrows mark descriptions matching the hits. The description includes the PDB number and chain designation (Hit), a description of the PDB entry including related PDB entries (Name), the probability of the hit based on the Hidden Markov Model (Probability), the probability of the match in an unrelated database (E-value), score for the secondary structure prediction (SS), number of amino acids aligned (Col), and the total length of the target in PDB (Target Length). The documentation for HHpred considers Probability the most important criterion with positive hits meeting at least one of these criteria: having a score >95% or having a score >50% and making reasonable biological sense [19]. *T. cruzi* and *L. major* orthologues have the same predicted folds.

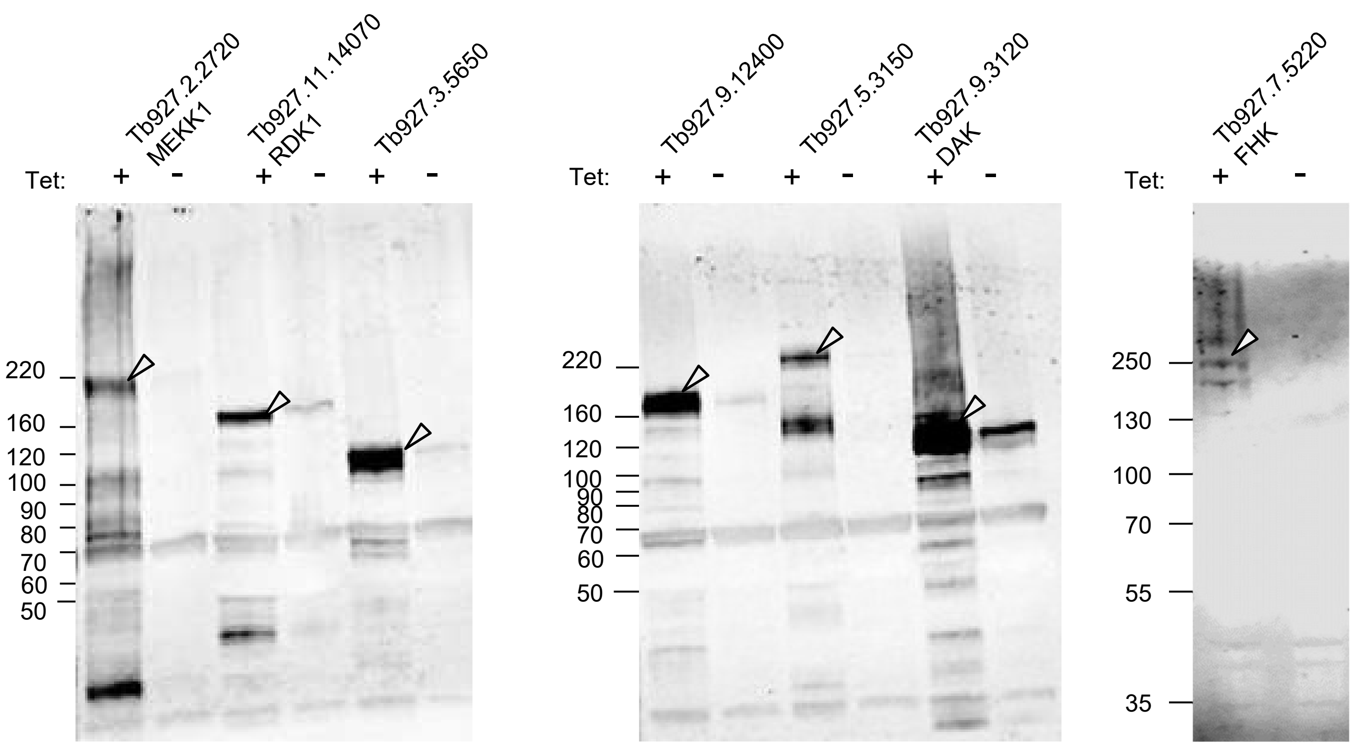


**Figure S1. Western blot showing expression of epitope-tagged PKs in *T. brucei* bloodstream forms.** Parasites were grown in the absence of Tet or in the presence of Tet to induce the V-5 tagged PK. The migration of each tagged protein is marked. The predicted molecular weights for each protein is provided in Table S1.
